# Kinesin light chain 1 (KLC1) interacts with NS1 and is a susceptibility factor for dengue virus infection in mosquito cells

**DOI:** 10.1101/2025.03.20.644413

**Authors:** Juan Manuel Castillo, Raymundo Cruz Pérez, Daniel Talamás, Juan E. Ludert

## Abstract

A hallmark of the dengue virus (DENV) infection is the manipulation of host cell membranes, lipid trafficking and lipid droplets (LDs), all cellular functions that depend on the cytoskeleton and the cytoplasmatic streaming system. We previously reported the interaction between DENV NS1 protein and members of the kinesin motor complex in the *Aedes albopictus* cell line C6/36. In this work, we present evidence indicating that the protein kinesin light chain 1 (KLC1) is indeed a susceptibility factor for DENV replicative cycle in mosquito cells. The interaction between NS1 and KLC1 was confirmed by proximity ligation and co-immunoprecipitation assays in cells harvested 24 hpi. In addition, transmission immunoelectron microscopy showed KLC1 decorating the surface of vacuoles in association with NS1. Increased levels of KLC1 were observed starting at 6 hpi, suggesting that virus infection stimulates KLC1 synthesis. Silencing KLC1 expression results in a reduction in viral genome synthesis, decreased secretion of NS1, and a reduction of virus progeny by nearly 1 log. In agreement, similar affectations were observed in infected cells transfected with a peptide that competes and interferes with the interaction between KLC1 and its cargo molecules. Of note, both silencing the expression or interfering with the function of KLC1 resulted in a disorganization of LDs, which decreased in number and increased in area, in mock or infected cells. These results, taken together, suggest that KLC1 is a host susceptibility factor for DENV in mosquito cells, necessary for the proper transport and homeostasis of LDs required for flavivirus replication. However, modest colocalization was observed between NS1 and LDs, and the significance of the KLC1 and NS1 interactions need to be further investigated.

## Introduction

The genus *Orthoflavivirus* represents a group of vector-borne viruses that have become a continuous threat to the world’s public health, with the potential to cause major outbreaks (1). Orthoflaviviruses like yellow fever virus (YFV), Japanese encephalitis virus (JEV), West Nile virus (WNV) are a major public health concern in their respective subtropical areas (2). In the past decade Zika virus (ZIKV) raised an epidemiological alert in the Latin America and the Caribbean regions, and nearly half of the earth’s population lives in dengue virus (DENV) risk areas (3). The highly mutagenic genome and the capacity of their vectors to adapt and take advantage of the world’s environmental shift make Orthoflaviviruses highly successful pathogens (4–6)

DENV is an icosahedral enveloped virus (7) with a +ssRNA genome of approximately 10 Kd that encodes for a polyprotein that is processed in 3 structural proteins (C, M, and E) and 7 non-structural proteins (NS1, NS2A, NS2B, NS3, NS4A, NS4B, and NS5) (8). In humans, virus mosquito inoculation is presumed to infect skin Langerhans dendritic cells, to then spread to lymph nodes, peripheral blood and spleen, where markers for replication have been observed (9,10). In *Ae. aegytpi* and *Ae. albopictus* mosquitoes, the first tissue to get infected after blood feeding is the midgut; the infection then spreads to the hemocoel and, finally the salivary glands(11).

A key feature in the *Orthoflavivirus* replication cycle is the manipulation of internal host membranes to form the replication organelles (RO) in endoplasmic reticulum (ER) derived membranes. Extensive remodeling of the ER membranes has been reported in both DENV and other closely related Orthoflaviviruses infected mammalian and mosquito cells (12–16). It has been proposed that these intricate structures not only serve as scaffolds, where viral replication and assembly occur, but they also favor enzymatic retention, substrate accumulation, and immune evasion in favor of viral replication (17–19). Therefore, several DENV proteins have acquired the capacity to interact with cellular membranes (14,20); for example, NS4A, which is thought to induce membrane curvature, anchoring itself in the cytosolic side of the RO via its transmembrane region (21), or NS1, which have been proposed to insert into the luminal side of membranes via its β-roll domain, also inducing membrane curvature and serving as scaffold for the replication complexes (RC). Others, like NS3, have been reported to redistribute and stimulate the fatty acid synthase to viral replication complexes to aid virus replication(22).

In addition, lipid hijacking is a well-recognized phenomenon in DENV infected cells (23,24). By altering lipid homeostasis, Orthoflaviviruses obtain the building blocks necessary to establish viral RC (14) and assembly of nascent viral particles (22,25). Moreover, processes like immune evasion and disease severity, which are involved in lipid metabolism, are related to the targeting of this system by DENV (26–28). Lipid droplets (LD) are organelles responsible for the storage and transport of neutral lipids, allowing the cell to have a reservoir of energy and structural building blocks for survival (29). The C protein of several Orthoflaviviruses is reported to interact with the surface of LD for nucleocapsid formation (30–32). Thus, targeting LD has been proposed as a potential strategy for the development of *Orthoflavivirus* antivirals (33).

The kinesin motor complex (KMC) is a system involved in the cytoplasmatic streaming of cargos through the microtubules in a highly coordinated and active pattern (34,35). It is composed of two heavy chains that perform the motor function and two light chains that serve as an adaptor that links the heavy chains with the cargo. Based on the subunits, the KMC can adopt several configurations, and this depends on the cell type and the cargo to be transported. The kinesin light chain 1 (KLC1), one of the subunits of the KMC is composed of three domains; the N-terminal domain connects with the heavy chains(36), the tetratricopeptide repeat domain (TPR) is the region that interacts with cargo proteins(37), and the C-terminal domain has been proposed to attach to membranes(38). Of note, the KMC is a system strongly related to LD motility (34) and there is evidence suggesting that KLC1 plays a significant role in the homeostasis of these organelles(39). Conversely, KLC1 and other members of the KMC have been reported as targets of viral effectors to favor viral replication (40–42), likely due to its ability to traverse all the cell and be involved in a myriad of cell processes. Recently, KLC1 was identified as part of the interactome of DENV-NS1 in mosquito C6/36 cells (43). Here, we present evidence indicating that KLC1 certainly interacts with NS1 and is a susceptibility factor for DENV in C6/36 cells, involved in the mobilization of neutral lipids via LDs. The results shed light on the mosquito cell-DENV interactions and identified potential new targets for antiviral strategies.

## Materials and Methods

### Cell cultures and viruses

The C6/36 cell line (CRL-1660; ATCC) was cultured with Minimum Essential Medium (MEM) (30-2003; ATCC) with 5% bovine fetal serum (FBS) and 100 U/ml penicillin-streptomycin (Gibco, 15140122) at 27°C and 5% of CO_2_. Confluent cell monolayers, grown in 6-well plates, were infected with DENV-2, strain New Guinea, at a MOI=3 in FBS free MEM, for 2 hours. Infections were allowed to proceed for 24 hpi, after replenishing the monolayers with MEM supplemented with FBS at 5% and antibiotics. BHK cells (BHK-21; ATCC CCL-10) were cultured in MEM with 5% of FBS and 100U/ml penicillin-streptomycin at 37°C and 5% of CO_2_.

### Cell viability

Cell viability was measured 24 h post-treatment with the MTS proliferation assay (Cell Titer 96 Aqueous; Promega, G3580) in 96-well plates, following the manufacturés recommendations.

### Gene silencing

Specific KLC1 siRNA duplexes were purchased from Integrated DNA Technologies and were targeted against the heptad repeat encoding region of *Aedes albopictus* KLC1 (>AALC636_019471.R27282). To carry out the silencing assays, C6/36 cell monolayers at 80% confluence, were transfected with 200nM of KLC1 siRNA duplex using 3 µL of lipofectamine 2000 (Invitrogen, 1168019), following the manufacturer instructions, and cells were processed for analysis 24 hpt. AllStars Negative Control siRNAS were used as control in the same concentration as the treatment (Qiagen, SI03650318). In the case of infected cells, siRNA transfection was done 2 hours post infection.

### Plaque formation

For progeny virus titration, cell supernatants were serially diluted (10-fold), and 0.2ml of each dilution was added in triplicate, onto confluent monolayers of BHK-21 cells grown in 6-well plates, in SFB free MEM for 2 hours at 37°C. Then, 1 mL of carboxymethyl cellulose was added and the cells were incubated for 5 days at 37°C and 5% CO_2_. Next, the carboxymethylcellulose was removed, and 500 µL of Naphthol Blue Black (Sigma-Aldrich, N9002) were added for 24 hours at room temperature. Finally, Naphthol Blue Black was removed, monolayers washed 3 times with tap water, and the plaques were counted. Results were expressed as PFU/ml.

### Indirect Immunofluorescence

At the appropriate times post infection, culture medium was removed, and monolayers fixed with 300 μl of a 4% paraformaldehyde solution for 30 minutes at 4°C. Then, cells were washed with 0.2% PBS-Tween 20 and permeabilized with 300 μl of a 0.2% PBS-Triton X-100 solution for 10 minutes at room temperature. Afterward, the glass slides were transferred to a humid chamber over a sheet of Parafilm® and 50 μl of blocking solution (Glycine 25 mg/ml, 1% FBS and 1% fish gelatin) were added for 2 hours at 37°C. The blocking solution was removed and 50 μl of the solution with primary antibodi es was added overnight at 4°C. The solution with antibodies was removed and the slides were washed 3 times for 10 minutes with PBS 0.2%-Tween 20, then 5 μl of a solution with secondary antibodies were added for 2 hours at 37°C. Finally, the solution with secondary antibodies was removed and a solution with DAPI (Sigma, D9542) 1:1000 was added for 10 minutes, and the slides were washed 3 times for 10 minutes with 0.2% PBS-Tween 20 and once for 5 minutes with Milli-Q water. The slides were mounted with 5μl of VectaShield (Vector, H-1000) and analyzed on a LSM 900 confocal microscope. For protein colocalization, approximately 100 cells were counted and the average colocalization index (Pearson correlation coefficient, PCC) was calculated with the Zeiss and Icy software.

### Proximity ligation assays

Proximity ligation assays were performed with a Duolink kit (Sigma-Aldrich, DUO92004) according to the manufacturer’s protocol. Briefly, infected cells were fixed at 24 hpi with a 4% paraformaldehyde solution for 30 minutes at 4°C. Then, the slides were transferred to a humidity chamber and one drop of the blocking solution was added for 1 hour at 37°C. Primary antibodies were added in a 1:100 dilution and incubated overnight at 4°C. Following this, primary antibodies were removed, and the slides washed with reagent A 2 times for 5 minutes each, followed by the addition of the minus and plus probes in a 1:5 dilution and incubated for 1 hour at 37°C. After incubation, the probes solution was removed and the slides were washed 2 times for 5 minutes each with reagent A; then, the ligase was added in a 1:40 dilution for 30 minutes at 37°C. After this, the ligase solution was removed and the slides were washed 2 times for 5 minutes each with reagent A. Afterwards, the ligase solution was removed, and the polymerase was added in a 1:80 dilution for 100 minutes at 37°C. Finally, the polymerase solution was removed, and the slides were washed 2 times for 10 minutes each with reagent B, then one time with reagent B 0.01X for 1 minute. Negative technical controls included mock cells exposed to both primary antibodies and infected cells exposed only to KLC1 antibody. Cells were analyzed under a confocal microscope Zeiss model LSM 900.

### Protein modelling and structural analysis

The 3D structure of C6/36 KLC1 (AALC636_019471.R27282) was modeled with Alpha Fold 3.0 (44). Molecular graphics and analyses were performed with UCSF ChimeraX (Resource for Biocomputing, Visualization, and Informatics at the University of California, San Francisco, with support from National Institutes of Health R01-GM129325, and the Office of Cyber Infrastructure and Computational Biology, National Institute of Allergy and Infectious Diseases(45).

### qRT-PCR

Genomic DENV RNA levels were determined by qRT-PCR. Total RNA was extracted from cell monolayers washed 3 times with PBS, with 500 µL of Trizol reagent (Thermo Fisher, 15596018) and 100 µL of chloroform (J.T Baker, 9180-02) were added for 3 minutes at room temperature. The solution was centrifuged at 12, 000xg for 15 minutes at 4°C. Supernatants were collected and 250 µL of isopropanol were added, the solution incubated at room temperature for 10 minutes and centrifuged at 12,000xg for 10 minutes at 4°C. Supernatants were discarded, and the pellet was resuspended with 1 mL of ethanol 75% and centrifuged at 7500xg for 5 min at 4°C. The supernatant was discarded, and the pellet was resuspended with 20µl of nuclease free water and treated with DNAse I (BioLabs Cat. M0303).

The qRT-PCRs were normalized with GAPDH mRNA, as a house keeping gene. For each reaction 5 µl of qPCR SyGreen 1-Step Go Hi-ROX (PB25.32-03, PCRBIOSYSTEMS), 0.5 µl of a stock of primers and 1 µl of RNA diluted in RNAse free water were added. Retro-transcription was performed at 50°C for 10 minutes, followed by enzyme inactivation and DNA denaturation at 95°C for 2 minutes; finally, 40 cycles of 5 seconds at 95°C and 30 seconds at 60°C were performed on an Eco Ilumina System. The threshold was adjusted with a non-infected sample and a non-template control. The results were analyzed with the EcoStudy software version 5.04890.

### Quantification of secreted NS1

The amount of cell-secreted NS1 was quantified using a homemade ELISA. Plates (Corning, 3590) were coated with 100 µl/well of a rabbit polyclonal NS1 antibody (5 µg final concentration) diluted in bicarbonate buffer (0.1M pH 9.6) overnight at 4°C. After removing the coating solution, wells were washed 4 times with PBS 0.05% Tween-20 and blocked with 200 µl of blocking solution (PBS, 1% BSA) for 1 hour at room temperature. After 4 washes with PBS 0.05% Tween-20, cell supernatants diluted in PBS 0.05% Tween-20, 0.1% BSA, were added and plates incubated for 2 h at room temperature. Wells were washed 4 times with PBS 0.05% Tween-20 and incubated with 100 µl of 7E11 biotinylated NS1 antibody diluted 1:4000 in dilution buffer (PBS 0.05% Tween-20,0.1% BSA) for 2 hours at room temperature. The wells were washed 4 times with PBS 0.05% Tween-20 and then 100 µl of streptavidin-HRP 1:8000 was added for 1 hour at room temperature. Finally, the wells were washed 4 times with PBS 0.05% Tween-20 and 100 µl of ABTS substrate (Sigma, A-1888) with H2O2 0.3% were incubated for 1 hour. A commercial recombinant DENV NS1 protein (R&D Systems) serially diluted was used as standard to build the calibration curve.

### Co-immunoprecipitation

The co-immunoprecipitation assays were performed using a commercial kit (Abcam, ab206996) according to the manufacturer’s instructions. For each sample, 25 µg of NS1 antibody were incubated in 500 µg of mock or infected cell lysates harvested at 24 hpi. Controls include mock and DENV infected cell lysates (input) and a mock infected eluted fraction (immunoprecipitate). Input and eluted samples were analyzed by western blot using antibodies against NS1 (Abcam, ab150111) and KLC1 (ABclonal, A5552) as primary antibodies. Results were documented with a Newton Mini FX7 Edge 18.11 - SN hardware.

### Immuno-electron microscopy

C6/36 cell monolayers were washed with PBS, scraped and centrifuged at 100xg for 8 minutes at 4°C. The pellet was resuspended with glutaraldehyde 0.5% and paraformaldehyde 4% and incubated at room temperature for 1 hour. Cells were washed with PBS and dehydrated with increasing concentrations of ethanol (50, 70, 90 and 100%). Afterwards, cells were embedded in LR-White pure resin (London Resin Co) and polymerized for 48 hours at 4°C in UV light. The thin sections were incubated overnight with anti-NS1 (Abcam, ab150111) and anti-KLC1 (ABclonal, A5552) antibodies at 4°C. The next day, samples were washed with PBS and incubated for 1 hour with goat anti-mouse (Ted Pella, Inc) and goat anti-rabbit IgG (H+L) (Ted Pella, Inc) secondary antibodies coupled with 15 and 30 nm gold particles, respectively. Finally, samples were washed with PBS and incubated with lead citrate and uranyl acetate and observed with a JEOL-LEM-011 transmission electron microscope.

### Lipid droplets staining and counting

For quantification of LD, cells were stained with Oil Red (Matheson Coleman & Bell) for 30 minutes; then, the slides were washed 3 times with PBS-Tween for 10 minutes each. Finally, LD were visualized using the indirect immunofluorescence protocol described above. LD were counted with the Image J software(46) with the Analyze particles plug -in. The threshold and detection range were set for the control images; once these parameters were established, they were applied for the analysis of the LD in the experimental samples.

### Peptide assays

KINTAG is a peptide of 17 residues with a FITC tag coupled by an Ahx linker (pepMic Co., Ltd, Jiangsu, China), that binds to the cargo region of KLC1(37). To evaluate the peptide transduction effect into the cell’s viability and LD homeostasis, C6/36 cell monolayers were washed once with PBS and incubated with KINTAG peptide, diluted in MEM-FBS 5% to a final concentration of 0.1 µg/µL, for 24 hours at 27 °C in a 5% CO_2_ atmosphere. The effect of the KINTAG on DENV infection was evaluated by treating the cells at 2 hpi, and the infection allowed to proceed until 24 hpi. At this time cells were harvested and processed for further analysis.

### Statistical analysis

Results were analyzed for statistical difference using Graph Pad Prism version 8.0.2.

## Results

### KLC1 interacts with NS1 in DENV infected C6/36 cells

Previous evidence from our group suggested interactions between KLC1 and NS1 in infected C6/36 cells (43). To corroborate this finding, immunofluorescence, PLA and immunoprecipitation assays were carried out in DENV infected C6/36 fixed at 24 hpi, using mock infected cells as controls (Figure 1). Colocalization between KLC1 and NS1 was observed, mainly in the cytoplasm (PCC=0.52) (Figure 1A). In addition, positive signals were observed in the cytoplasm of infected cells by PLA, while signal was observed in the negative controls, run in parallel, indicating specific interaction between KLC1 and NS1 (Figure 1B). Finally, pull-down of NS1 from DENV infected cell lysates showed the presence of one KLC1 isoform in the immunoprecipitated fraction, absent in mock-infected cell lysates (Figure 1C). All these results taken together indicate that KLC1 and DENV NS1 interact in C6/36 infected cells.

**Figure 1.**
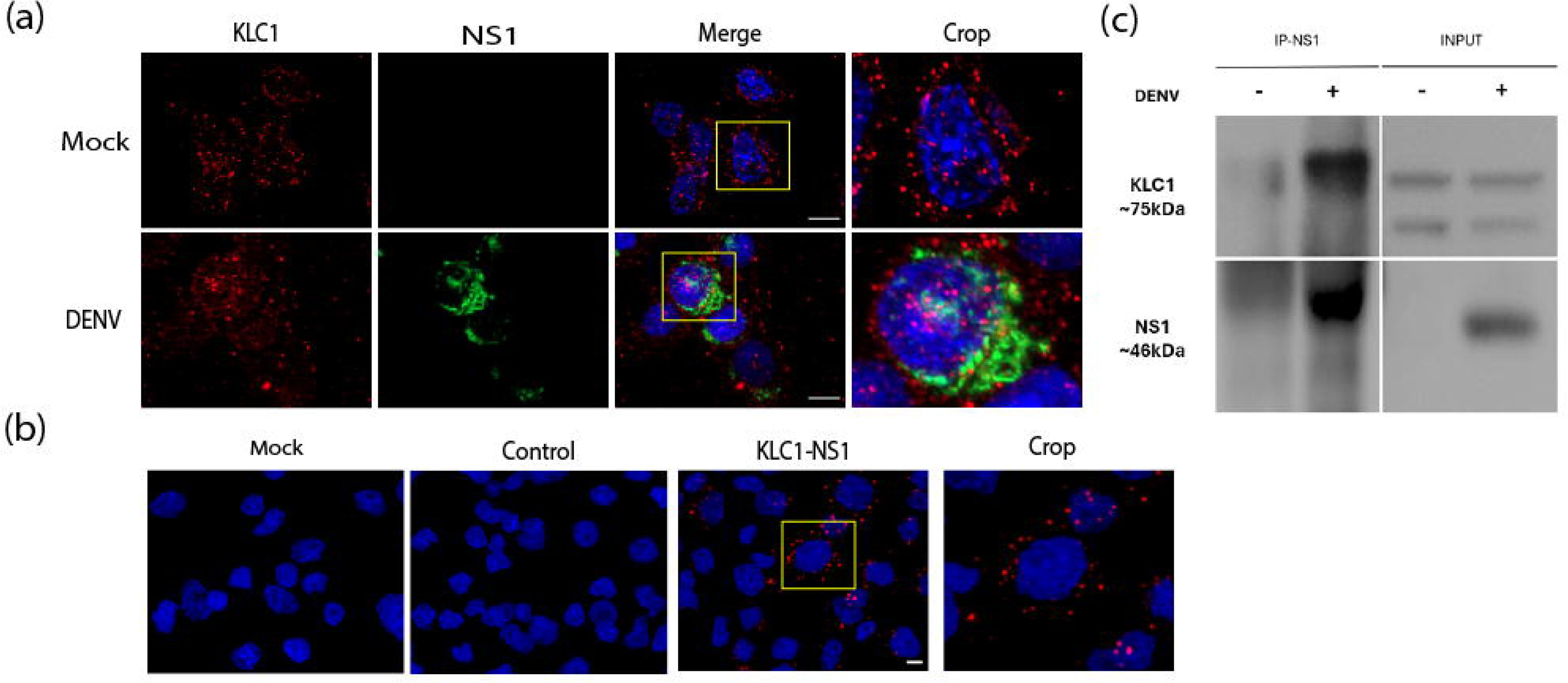
KLC1 interaction with DENV NS1 in infected C6/36 cells. **A)** Colocalization assays of KLC1 and DENV NS1 (Pearson = 0.52). **B)** Proximity ligation assays of KLC1 and DENV NS1. Mock infected cells treated with anti-KLC1 (*mock*) and infected cells with primary anti-KLC1 antibody omitted (*control*) were included as negative controls*. Crop in A and B;* enlargement of the cells enclosed in the yellow square. **C)** Immunoprecipitation assays. Cell lysates were immunoprecipitated with anti-NS1 antibodies bound to Sepharose beads. The presence of DENV KLC1 in the immunoprecipitant was revealed by Western blot. Changes in protein migration observed in the immunoprecipitate are due to elution buffer composition, and were consistently observed in experimental replicas. In all cases, mock and infected cells were harvested at 24 hpi. C6/36 average cell diameter= 12.5 µm. Bar= 5 µm. Representative images of at least 3 independent experiments shown.

### KLC1 modeling and docking assays with NS1

To gain insights into the KLC1 and NS1 interactions, the full KLC1 molecule was modeled via AlphaFold 3.0, since only the TPR domain crystallographic structure is available(37). The KLC1 model obtained showed a pTM of 0.85, indicating high confidence in the predicted structure (Figure 2A). Moreover, comparison of the TPR region of *Homo sapiens* with the predicted TPR region of C6/36 KLC1 showed a RMSD value of 0.53Å (Figure 2B), indicating high structural resemblance. Finally, docking assays between the TRP domain of KLC1 and the NS1 dimer showed a possible interaction between the β-roll region of NS1 dimer and the inner groove of the TPR domain with a pTM of 0.68 and an ipTM of 0.63 (Figure 2C).

**Figure 2.**
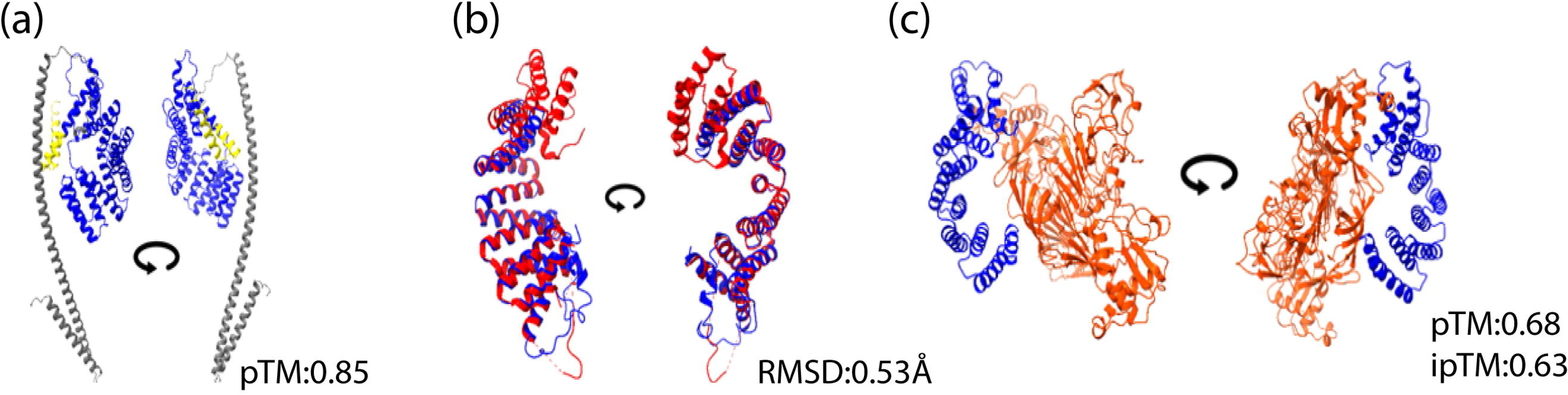
In silico modeling of mosquito KLC1. **A)** Alpha-fold 3.0 modeling of the KLC1 of *Aedes albopictus*. TRP domain shown in blue. **B)** Superimposition of the TPR cargo binding domain of mosquito (blue) and human (red) KLC1. **C)** Modeled interaction of the TPR (blue) and NS1 dimers (orange). pTM: predicted template modeling. ipTM: Interface predicted template modeling. RMSD: root mean square deviation. Circular arrow=180° image rotation.

### KLC1 colocalizes with NS1 in the surface of membrane bound organelles

To gain further information on the interaction between KLC1 and NS1, infected cells were analyzed by immuno-electron microscopy to determine the sub-cellular regions where these interactions take place. In mock cells, the mark for KLC1 was observed on the surface of membrane bound organelles of different sizes (Figure 3). In infected cells, the mark for KLC1 remained on the surface of these organelles but was accompanied by the mark of NS1 (Figure 3). No mark for NS1 was observed in the mock infected cells. These results suggest that the interaction between KLC1-NS1 occurs in the surface of membrane bound organelles, and in agreement with the capacity of both KLC1 and NS1 to interact with membranes(38,47,48).

**Figure 3.**
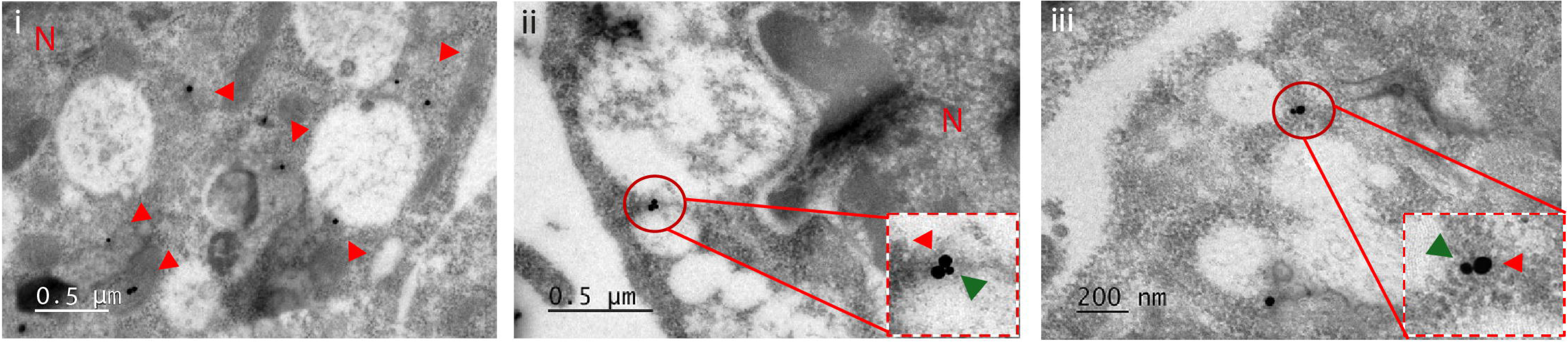
Immuno-electron microscopy of C6/36 cells infected with dengue virus. KLC1 was labelled with 30nm (red arrowheads), and NS1 with 15 nm (green arrow heads) gold particles. **i)** Mock infected cells; **ii)** and **iii)** infected cells. Cells were fixed at 24 hpi. KLC1 and NS1 were found to colocalized around vacuoles-like structures. N= cell nuclei.

### KLC1 is over-expressed in DENV infected C6/36 cells

One common signature of *Orthoflavivirus* replication is the overexpression of host susceptibility cellular proteins, which aid viral replication (43,49–52). In this sense, significantly increased levels of KLC1 were observed by western blot, in infected cells harvested at 24 hpi (Figure 4), when compared to mock infected cells. Increments in KLC1 levels were observed starting at 6 hpi, but did not reach significance until 12 hpi (data not shown). These results suggest that KLC1 is a host factor required by DENV in mosquito cells.

**Figure 4.**
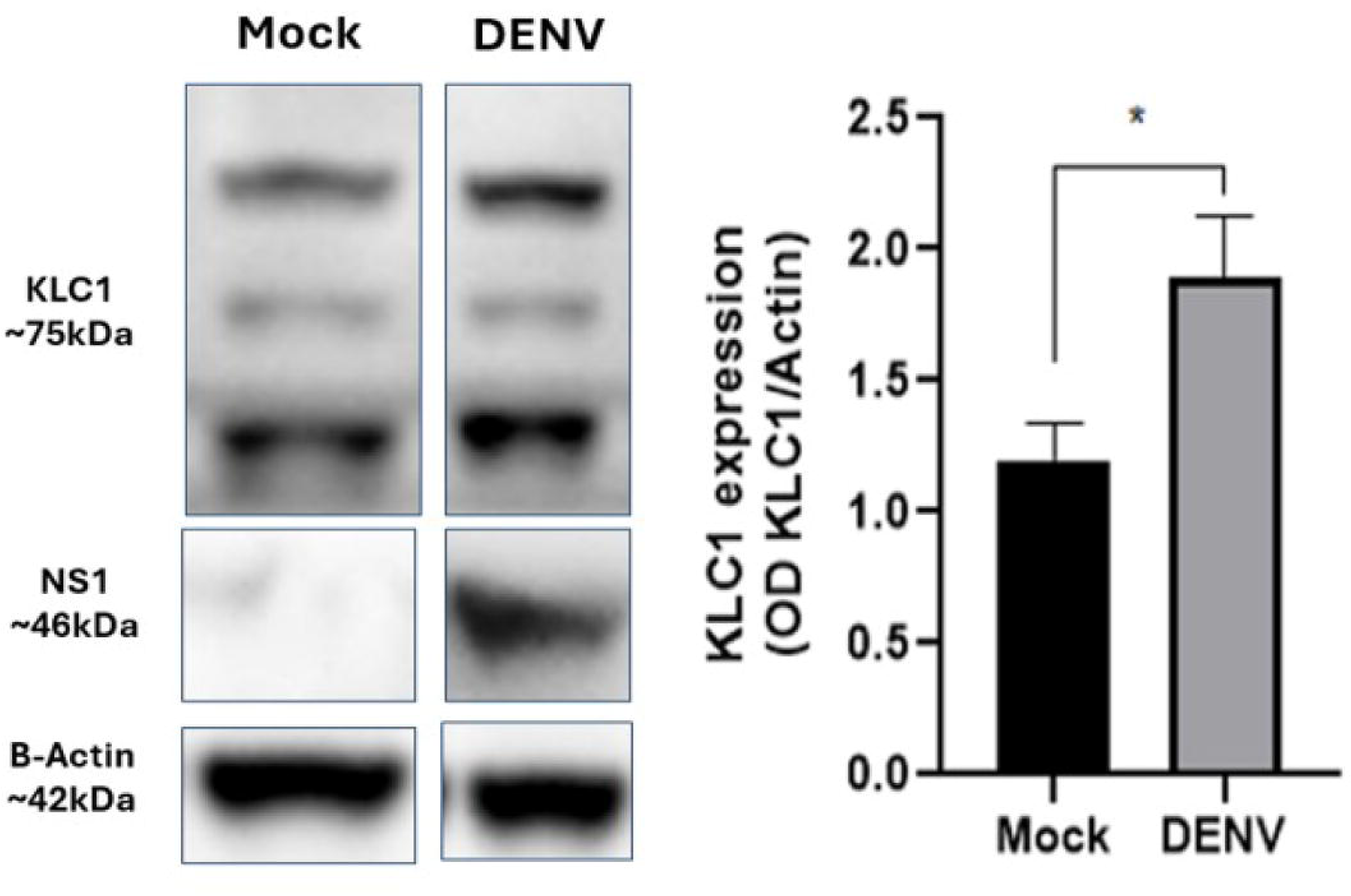
KLC1 expression levels in dengue virus infected cells. Cell lysates of mock and DENV infected cells were analyzed by western blot. Relative KLC1 protein levels are expressed as a ratio to β-actin used as load control. Representative images of 3 independent experiments are shown. * p≤0.005.

### KLC1 is a susceptibility factor for DENV infection in C6/36 cells

To gain insight into the participation of KLC1 in the replicative cycle of DENV, the expression of KLC1 was knocked down using siRNAs. A reduction of 50% in the expression of KLC1, as assayed by western blot, was achieved in mock-infected cells transfected with an effective siRNA-KLC1 concentration of 200 nm (Figure 5A); a concentration proven to be non-toxic for C6/36 cells (Supplemental Figure 1). The transfection of infected C6/36 cells with siRNA-KLC1 resulted in a ∼50% reduction in the levels of the NS1 protein, measured at 24 hpi (Figure 5B). The amount of secreted, soluble NS1 was also reduced by 50% (Figure 5C), compared to irrelevant siRNA transfected cells, reflecting the decrease observed in the intracellular levels of NS1. These results suggested that KLC1 is necessary for NS1 protein synthesis. In addition, viral genome synthesis and virus progeny were also evaluated. As shown in Figure 5D and E, a significant reduction in the relative expression of genomic RNA and in virus yield were observed in KLC1 silenced cells, in comparison with cells transfected with the irrelevant siRNA used as control. These results suggest that KLC1 silencing affects the overall DENV replication cycle.

**Figure 5.**
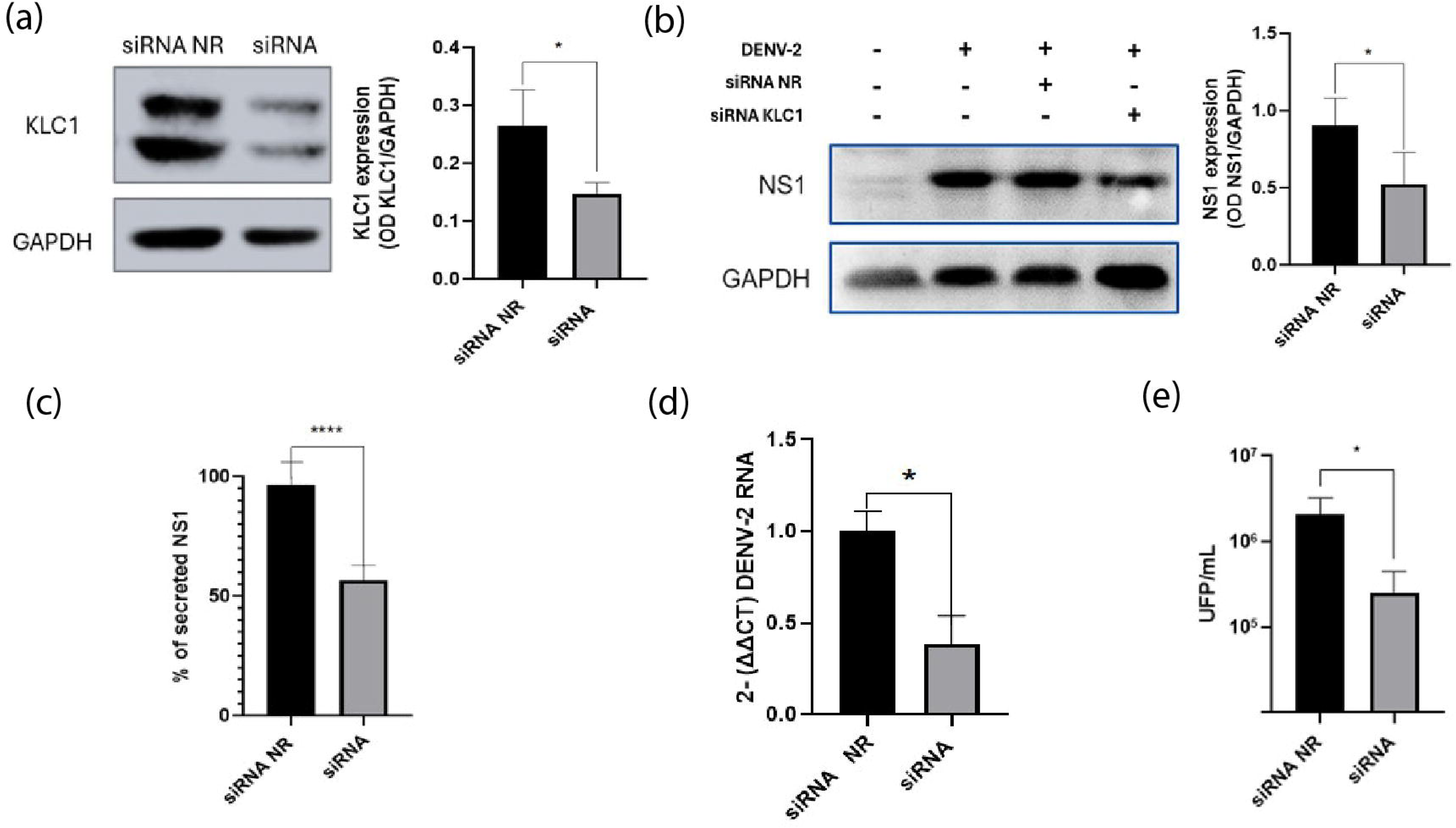
Effect on dengue virus replicative cycle of silencing the expression of KLC1. **A)** Transfection of C6/36 cells with siRNA specific for KLC1 or a non-related (NR) siRNA. Cells were lysed 24 hpt and the level of protein expression was analyzed by western blot. Relative KLC1 protein levels are expressed as a ratio to GADPH. **B)** Expression levels of NS1 in dengue virus infected cells silenced or not for KLC1 and lysed 24 hpi. NS1 levels are expressed as a ratio to GADPH. **C)** Amount of secreted NS1 in cells silenced or not for the expression of KLC1. Secreted NS1 was measured by an in-house ELISA, an expressed as percentage of the control condition, taken as 100%. **D)** Virus genome levels. RNA levels were determined by qRT-PCR. **E)** Virus yield. Virus progeny was determined by plaque assay. Cells were lysed and supernantants collected at 24 hpi. * p≤0.005. **** p≤0.001. n≥3.

Thus, confocal microscopy assays staining for NS1 and NS4A were carried out as a measure of the integrity of the replication complexes, given that NS4A is an integral part of these structures (21). A reduction in both NS1 and NS4A signals was observed in infected cells silenced for KLC1 in comparison with non-related siRNA transfected cells (Figure 6). Moreover, the colocalization between NS1 and NS4A was reduced in the KLC1 silenced cells (PCC 0.40 vs 0.31), suggesting affectation in the integrity of the replication complexes. These results all together revealed that KLC1 is necessary for the DENV replication cycle in mosquito cells.

**Figure 6.**
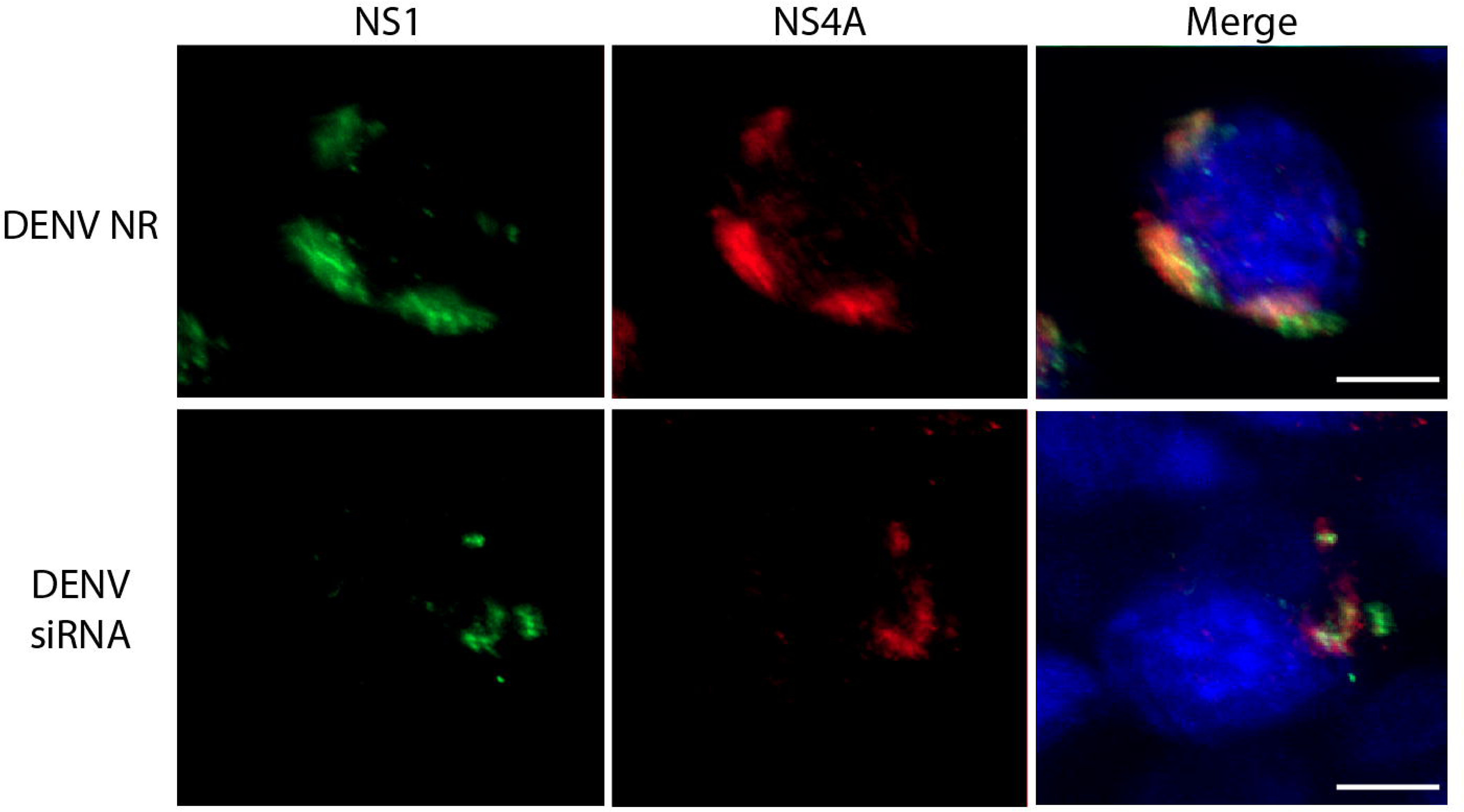
Effect on dengue virus replication complexes of cells silenced for KLC1. **A)** Cells transfected with a non-related (NR) siRNA. **B)** Cells transfected with a KLC1 specific siRNA. Cells were fixed 24 hpi. Bar= 5 µm. Representative images of 3 independent experiments are shown.

### The KINTAG peptide affects DENV replicative cycle

To corroborate the results obtained in cells silenced for KLC1, infected cells were transduced with KINTAG, a 17-residue peptide with high affinity to the TPR region of KLC1 of mice(37) . The rationale for KINTAG use is to interfere with the ability of the TPR region of KLC1 to bind cargo proteins. The peptide was used at a final concentration of 0.1µg/µL, the highest non-lethal concentration denoted for C6/36 cells (Supplemental Figure 1). Figure 7A shows the cell internalization of KINTAG evaluated at 24 h after exposure by confocal microscopy. Infected cells treated with KINTAG 2 hpi, showed significantly reduced levels of NS1 protein synthesis and secretion, and also significantly reduced viral genome and virus yield levels, at 24 hpi (Figure 7B-E). These results corroborate the results obtained using siRNAs and support the notion that a functional KLC1 is a necessary factor for the DENV replication cycle. Finally, docking assays indicated that, as expected, KINTAG has the potential to interact with the TPR region of C6/36 KLC1, showing that the association between the peptide and the inner groove of the TPR region is similar to the interaction observed with the TRP crystal 6SWU (37)(Figure 7F).

**Figure 7.**
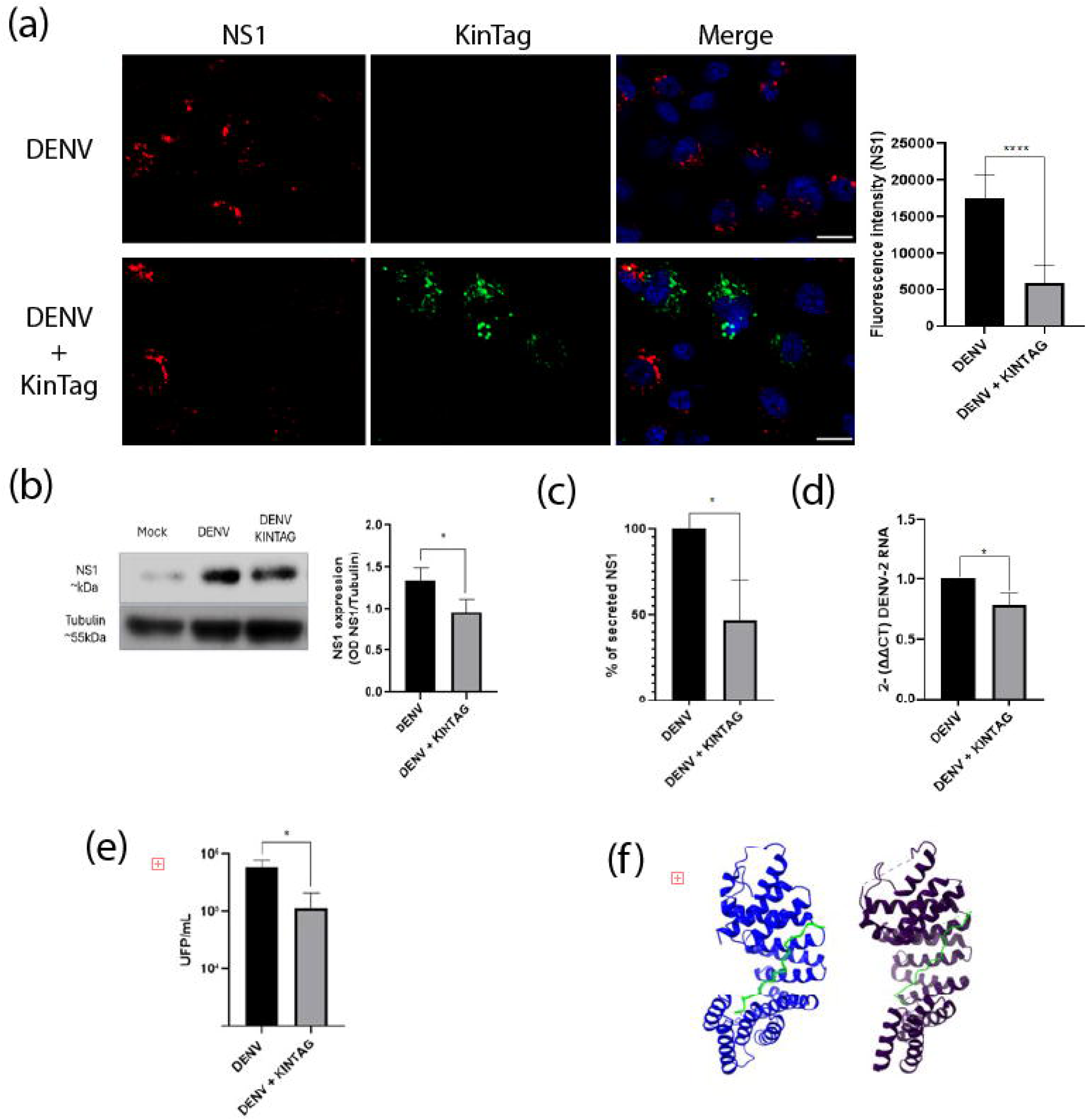
Effect of the KINTAG peptide in the replication of DENV. **A)** Fluorescence intensity of NS1 in cells transfected with KINTAG peptide fixed 24hpi. **B)** Expression levels of NS1 in DENV infected cells with KINTAG peptide 24hpi. NS1 levels are expressed as a ratio to tubulin. **C)** Amount of secreted NS1 in cells with KINTAG peptide. Secreted NS1 was measured by an in-house ELISA, an expressed as percentage of the control condition, taken as 100%. **D)** Virus genome levels. RNA levels were determined by qRT-PCR. **E)** Virus yield. Virus progeny was determined by plaque assay. Cells were lysed and supernatants collected at 24 hpi. **F)** Comparison between 6SWU crystal structure (purple) and C6/36’s KLC1 TPR and KINTAG peptide interaction predicted by Alpha Fold (Blue), KINTAG peptide is shown in green. Bar= 10µm. * p≤0.005. **** p≤0.001. n≥3.

### KLC1 is involved in lipid droplets homeostasis

The KMC is known to be involved in the trafficking and mobilization of LD, organelles that are required for the DENV replication cycle(53,54). Interestingly, major changes in the number or area of LD were observed in either mock or infected C6/36 cells treated with siRNA-KLC1 (Figure 8A and 8B). In cells with KLC1 synthesis diminished, the number of LD per cell was significantly reduced, while the mean area of LD increased (Figure 8C), suggesting that either coalescence of LD is taking place or proper dispersion is affected. Interestingly, infected cells with altered LD patterns also presented a marked reduction in NS1 fluorescence signal (Figure 8B). To further evaluate a role for KLC1 in LDs homeostasis, mock-infected cells were transfected with KINTAG. A significant reduction in the number of LDs per cell was also observed in the treated cells (Figures 8D and 8E). In agreement with the literature (39), these results indicate a role for KLC1 in the homeostasis of LD, thus hinting at a mechanism for the effects of KLC1 silencing or function alterations on the DENV replication cycle. Interestingly, only modest colocalization between LD and NS1 (PCC = 0.34) was observed, and in a pattern that indicates that only a fraction of both pools is adjacent to each other (Supplemental Figure 1C). Thus, the full significance of the interaction of NS1 with KLC1 needs further investigation.

**Figure 8.**
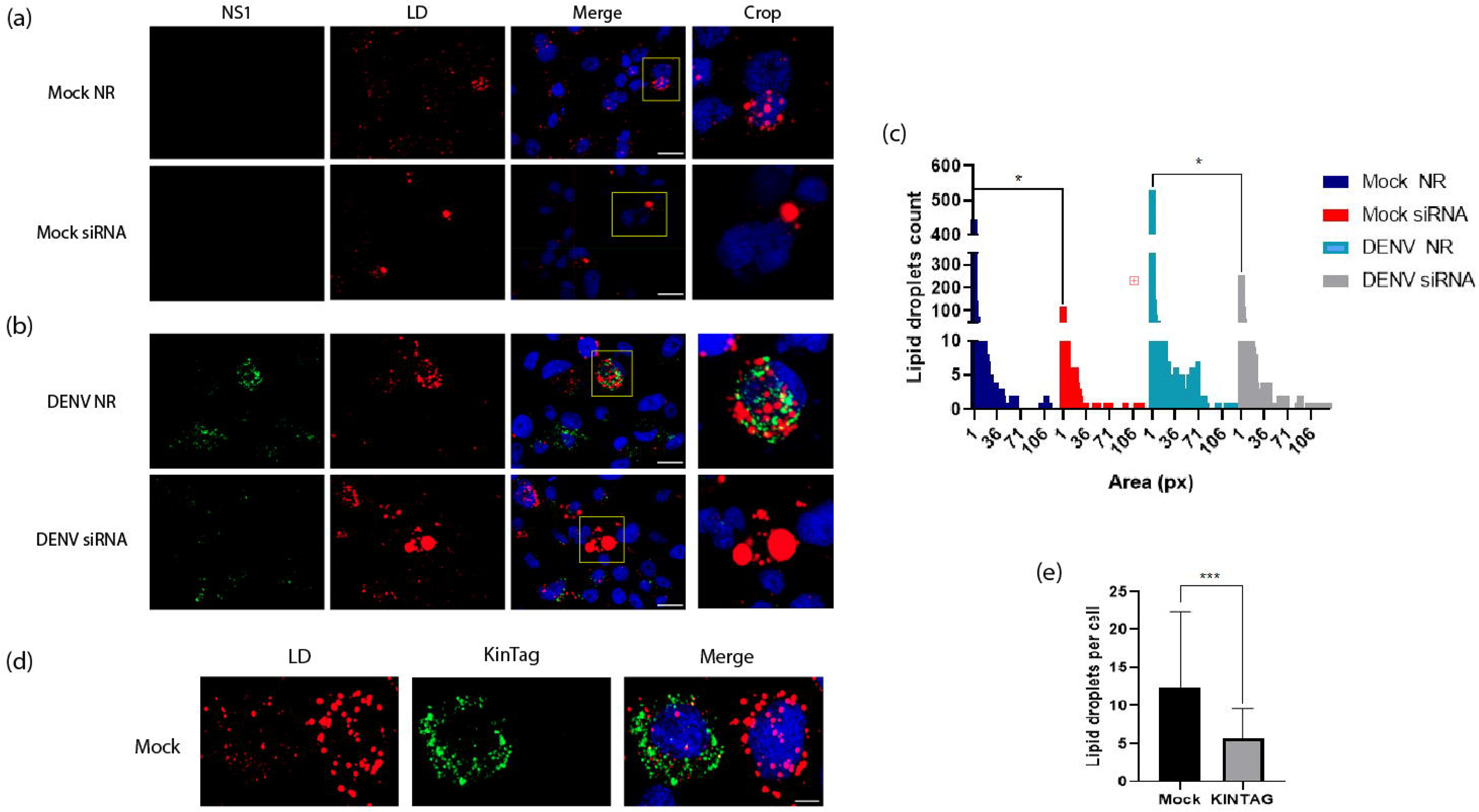
Effect of KLC1 silencing or function disruption on lipid droplets (LD) homeostasis. **A)** Mock infected cells silenced (siRNA) or not (NR) for KLC1 expression. **B)** DENV infected cells silenced or not for KLC1 expression. Cells were fixed at 24 hpi. **C)** Lipid droplet count in mock and infected cells silenced or not for KLC1. The graph represents the LD count and area measured in pixels of 130 cells for each experimental condition. **D)** Mock-infected cells transfected with KINTAG, fixed at 24 hpt. **E)** Lipid droplet count in KINTAG transfected cells. Lipid droplets were stained with oil red-O. Bar= 10µm. Representative images of 3 independent experiments are shown. * p≤0.005.

## Discussion

The effective control of dengue will require a better understanding of the interactions between the DENV and the mosquito vector. Yet, aspects like virus-cell interactions or virus-host protein interactions are better understood for the mammalian host than for the vector mosquito, a knowledge gap that needs to be closed. An interactome generated using a proximity biotinylation system and mass spectrometry revealed KLC1 as part of the interactome of NS1 in C6/36 cells C(43). Interestingly, KLC1 was not detected in a previous DENV NS1 interactome based on 3 different mammalian cell lines(55). These dissimilar observations suggested that KLC1 may be a susceptibility factor for DENV replicative cycle in mosquitoes. In this work, the interaction between DENV NS1 and KLC1 in C6/36 cells was demonstrated by 3 different specific assays, thus corroborating our previous identification of KLC1 as part of the interactome map of NS1 in C6/36 cells (43).

A role of KLC1 in the anterograde transport of viral protein cargos has been proposed in the context of several DNA virus infections (38,40,56). Immunoelectron microcopy micrographs suggest that KLC1-NS1 interactions take place in the cytoplasm around vacuoles-like, membrane-bound organelles. Both KLC1 and NS1 have the capacity to interact with lipid membranes and induce their bending (38,47,48). KLC1 is a protein proposed to interact with membrane-bound organelles by its C-terminal domain and to interact with cargos through the TPR domain located in the cytoplasm; meanwhile, NS1 is mainly located in the ER luminal side (38,47,48). Thus, the interaction between KLC1 and NS1 reported in this study could happen through ER or other internal membranes. Alternatively, NS1 can interact directly in the cytoplasm with the TPR region of KLC1, making NS1 a cargo protein. This possibility is supported by the docking assays between NS1 and the C6/36 KLC1 TPR region showing a possible interaction between the TPR domain of KLC1 and the β-roll domain of NS1 (Fig. 2C). Of note, evidence suggesting the presence of a fraction of NS1 in the cell cytoplasm is indirect but has been accumulating in recent years (43,55,57).

Regardless of the KLC1 cargo condition of NS1, our results indicate that KLC1 is a susceptibility factor for DENV in mosquito cells. RNA replication, protein synthesis, NS1 secretion, and virus progeny were all processes significantly affected in infected cells silenced for the expression of KLC1 or transduced with a high affinity peptide for the TRP domain. In addition, the levels of KLC1 were found to significantly increase at 24 hpi, in agreement with previous results obtained measuring KLC1 RNA levels in DENV infected C6/36 cells (58); increased expression of a cell susceptibility factors has been frequently observed in Orthoflaviviruses infected cell (49–51). A plausible explanation for the overall “up-river” effect observed in the absence of KCL1 is that the KMC participates in the reorganization and formation of the RO in the infected cells. Indeed, in the absence of KLC1, the colocalization coefficient index between NS1 and NS4A was significantly reduced, suggesting a disruption of the RC. These results are in line with a series of studies that address the hijacking of the cytoskeleton and the cytoplasmatic streaming system by Orthoflaviviruses (59–62). Noteworthy, KLC1 has been studied in the context of other viral infections (42,56) and was proposed to be involved in the anterograde transport of viral cargoes, positioning it as a frequent target of viral effectors.

The identification of the membrane-bound organelles where KLC1-NS1 interaction takes place, as suggested by the immunoelectron microscopy assays, was hindered by the micrograph’s resolution, but these structures resembled LD’s and vesicle-like structures (63–65). LDs have been recognized as required for DENV replication and proposed as antiviral targets (33). Several subunits of the KMC, including KLC1, have been reported to be involved in the mobilization of LD through the microtubules (34,39). In this regard, we found significant colocalization between the base of the microtubules and the NS4A protein (Supplementary Figure 2B), an integral component of the RCs, suggesting a connection between these two structures as observed in other Orthoflaviviruses (62,66). LD’s and replicative organelles alike originate from the membranes of the ER (14,67,68), and have been observed to be adjacent to each other in some Orthoflaviviruses infections (69,70). When silencing KLC1 or transducing KINTAG peptide, we observed significant alterations in LDs homeostasis, including a drop of LD’s count per cell, and the appearance of LD’s of augmented area (Figure 8), indicating that in the absence of KLC1, the transit of neutral lipids was disturbed. Neutral lipids are required for β-oxidation, ATP production, and membrane remodeling, which are all critical processes for DENV replication cycle (28,71–73). In support, silencing other members of the KMC like KIF5-B(35)causes alterations on LDs homeostasis similar to the ones observed in this study. The possible interaction between NS1 and LDs was directly evaluated by confocal microscopy, and modest colocalization was observed, suggesting that only a fraction of NS1 pools interact with LDs (Supplementary Figure 2A). These observations contrast with results reported for the C protein, where the entire surface of LD’s is decorated with the viral protein as part of the nucleocapsid formation processes (53).

In summary, this study reveals a role for KLC1 as a susceptibility factor for DENV replication in mosquito cells. The participation of KLC1 as part of the KMC in the lipid hijacking processes in mosquito cells by DENV, particularly in relation to LDs, appears to be one of the functions that KLC1 fulfills during the replication cycle. Nonetheless, other KLC1 functions were not evaluated in relation to the DENV replicative cycle, and the mechanistic significance of the NS1-KLC1 interaction was not fully elucidated. Despite these limitations, this work contributed to the understanding of virus-cell interactions and uncovered new potential antiviral targets for Orthoflaviviruses in mosquito cells.

## Acknowledgments

We would like to acknowledge Drs. Ana Alcalá and Ana Lorena Gutiérrez for their critical reading of the manuscript.

## Figure legends

**Supplemental figure 1.** Effect of KLC1 silencing and KINTAG peptide transduction on C6/36 cells viability. A) KLC1 siRNA 200nM. B) KINTAG peptide in concentrations ranging from 0.12µg/µl to 1.0 µg/µl. MTS assay was performed 24 hpt. Results are the means of 3 independent experiments. ** p≤0.01.

**Supplemental figure 2.** A) Colocalization between LD’s and NS1; Pearson correlation coefficient=0.34 B) Colocalization between Tubulin and NS4A; Pearson correlation coefficient=0.72. DENV infected C6/36 were fixed at 24hpi. Colocalization areas are shown in yellow and anti-colocalization areas are shown in purple. Bar= 5 µm.

## References

1. Pierson TC, Diamond MS. The continued threat of emerging flaviviruses. Vol. 5, Nature Microbiology. Nature Research; 2020. p. 796–812.

2. Huang YJS, Higgs S, Horne KME, Vanlandingham DL. Flavivirus-Mosquito interactions. Vol. 6, Viruses. MDPI AG; 2014. p. 4703–30.

3. Messina JP, Brady OJ, Golding N, Kraemer MUG, Wint GRW, Ray SE, et al. The current and future global distribution and population at risk of dengue. Nat Microbiol. 2019 Sep 1;4(9):1508–15.

4. Enukoha C, Talbalaghi A, Hassandoust S, Fossati F, Bazzoni M, Parisato S, et al. Interplay of climate change with physiological changes in adult Aedes albopictus. Acta Trop. 2024 Dec 1;260.

5. Dolan PT, Whitfield ZJ, Andino R. Mechanisms and Concepts in RNA Virus Population Dynamics and Evolution. Annual Reviews [Internet]. 2018;59:25. Available from: 10.1146/annurev-virology-

6. Egizi A, Fefferman NH, Fonseca DM. Evidence that implicit assumptions of ‘no evolution’ of disease vectors in changing environments can be violated on a rapid timescale. Philosophical Transactions of the Royal Society B: Biological Sciences. 2015;370(1665):1–10.

7. Rey FA, Stiasny K, Vaney M, Dellarole M, Heinz FX. The bright and the dark side of human antibody responses to flaviviruses: lessons for vaccine design. EMBO Rep. 2018 Feb;19(2):206–24.

8. Khan MB, Yang ZS, Lin CY, Hsu MC, Urbina AN, Assavalapsakul W, et al. Dengue overview: An updated systemic review. Vol. 16, Journal of Infection and Public Health. Elsevier Ltd; 2023. p. 1625–42.

9. Begum F, Das S, Mukherjee D, Mal S, Ray U. Insight into the tropism of dengue virus in humans. Vol. 11, Viruses. MDPI AG; 2019.

10. Reyes-del Valle J, Salas-Benito J, Soto-Acosta R, del Angel RM. Dengue Virus Cellular Receptors and Tropism. Vol. 1, Current Tropical Medicine Reports. Springer Verlag; 2014. p. 36–43.

11. Salazar MI, Richardson JH, Sánchez-Vargas I, Olson KE, Beaty BJ. Dengue virus type 2: Replication and tropisms in orally infected Aedes aegypti mosquitoes. BMC Microbiol. 2007;7.

12. Wong HH, Crudgington DRK, Siu L, Sanyal S. Flaviviruses induce ER-specific remodelling of protein synthesis. PLoS Pathog. 2024 Dec 1;20(12).

13. Ci Y, Liu ZY, Zhang NN, Niu Y, Yang Y, Xu C, et al. Zika NS1-induced ER remodeling is essential for viral replication. Journal of Cell Biology. 2020 Feb 3;219(2).

14. Welsch S, Miller S, Romero-Brey I, Merz A, Bleck CKE, Walther P, et al. Composition and Three-Dimensional Architecture of the Dengue Virus Replication and Assembly Sites. Cell Host Microbe. 2009 Apr 23;5(4):365–75.

15. Ko KK, Igarashi A, Fukai K. Electron Microscopic Observations on Aedes albopictus Cells Infected With Dengue Viruses. Vol. 62, Archives of Virology. 1979 May.

16. Grief C, Galler R, Co Ãrtes LMC, Barth OM. Intracellular localisation of dengue-2 RNA in mosquito cell culture using electron microscopic in situ hybridisation. Vol. 142, Arch Virol. 1997.

17. Hsu NY, Ilnytska O, Belov G, Santiana M, Chen YH, Takvorian PM, et al. Viral reorganization of the secretory pathway generates distinct organelles for RNA replication. Cell. 2010;141(5):799–811.

18. Reiss S, Rebhan I, Backes P, Romero-Brey I, Erfle H, Matula P, et al. Recruitment and activation of a lipid kinase by hepatitis C virus NS5A is essential for integrity of the membranous replication compartment. Cell Host Microbe. 2011 Jan 20;9(1):32–45.

19. Marceau CD, Puschnik AS, Majzoub K, Ooi YS, Brewer SM, Fuchs G, et al. Genetic dissection of Flaviviridae host factors through genome-scale CRISPR screens. Nature. 2016 Jul 6;535(7610):159–63.

20. Morita E, Suzuki Y. Membrane-associated flavivirus replication complex—its organization and regulation. Vol. 13, Viruses. MDPI AG; 2021.

21. Miller S, Kastner S, Krijnse-Locker J, Bühler S, Bartenschlager R. The non-structural protein 4A of dengue virus is an integral membrane protein inducing membrane alterations in a 2K-regulated manner. Journal of Biological Chemistry. 2007;282(12).

22. Heaton NS, Perera R, Berger KL, Khadka S, LaCount DJ, Kuhn RJ, et al. Dengue virus nonstructural protein 3 redistributes fatty acid synthase to sites of viral replication and increases cellular fatty acid synthesis. Proc Natl Acad Sci U S A. 2010;107(40).

23. Melo CFOR, Delafiori J, Dabaja MZ, de Oliveira DN, Guerreiro TM, Colombo TE, et al. The role of lipids in the inception, maintenance and complications of dengue virus infection. Sci Rep. 2018 Dec 1;8(1).

24. Hsia JZ, Liu D, Haynes LP, Cruz-Cosme R, Tang Q. Lipid Droplets: Formation, Degradation, and Their Role in Cellular Responses to Flavivirus Infections. Vol. 12, Microorganisms. Multidisciplinary Digital Publishing Institute (MDPI); 2024.

25. Chang KS, Jiang J, Cai Z, Luo G. Human Apolipoprotein E Is Required for Infectivity and Production of Hepatitis C Virus in Cell Culture. J Virol. 2007 Dec 15;81(24):13783–93.

26. Barletta ABF, Alves LR, Nascimento Silva MCL, Sim S, Dimopoulos G, Liechocki S, et al. Emerging role of lipid droplets in Aedes aegypti immune response against bacteria and Dengue virus. Sci Rep. 2016 Feb 18;6.

27. Coelho DR, Carneiro PH, Mendes-Monteiro L, Conde JN, Andrade I, Cao T, et al. ApoA1 Neutralizes Proinflammatory Effects of Dengue Virus NS1 Protein and Modulates Viral Immune Evasion. J Virol [Internet]. 2021;95(13):1974–94. Available from: 10.1128/JVI

28. Yousefi M, Lee WS, Yan B, Cui L, Yong CL, Yap X, et al. TMEM41B and VMP1 modulate cellular lipid and energy metabolism for facilitating dengue virus infection. PLoS Pathog. 2022 Aug 1;18(8).

29. Fujimoto T, Parton RG. Not just fat: The structure and function of the lipid droplet. Cold Spring Harb Perspect Biol. 2011 Mar;3(3):1–17.

30. Iglesias NG, Mondotte JA, Byk LA, De Maio FA, Samsa MM, Alvarez C, et al. Dengue Virus Uses a Non-Canonical Function of the Host GBF1-Arf-COPI System for Capsid Protein Accumulation on Lipid Droplets. Traffic. 2015 Sep 1;16(9):962–77.

31. Shang Z, Song H, Shi Y, Qi J, Gao GF. Crystal Structure of the Capsid Protein from Zika Virus. J Mol Biol. 2018 Mar 30;430(7):948–62.

32. Martins AS, Carvalho FA, Nascimento AR, Silva NM, Rebelo T V., Faustino AF, et al. Zika virus capsid protein closed structure modulates binding to host lipid systems. Protein Science. 2024 Sep 1;33(9).

33. Hyrina A, Meng F, McArthur SJ, Eivemark S, Nabi IR, Jean F. Human Subtilisin Kexin Isozyme-1 (SKI-1)/Site-1 Protease (S1P) regulates cytoplasmic lipid droplet abundance: A potential target for indirect-acting anti-dengue virus agents. PLoS One. 2017;12(3).

34. Kilwein MD, Welte MA. Lipid Droplet Motility and Organelle Contacts. Vol. 2, Contact. SAGE Publications Inc.; 2019.

35. Rai P, Kumar M, Sharma G, Barak P, Das S, Kamat SS, et al. Kinesin-dependent mechanism for controlling triglyceride secretion from the liver. Proc Natl Acad Sci U S A. 2017;114(49).

36. Gauger AK, Goldstein LSB. The Drosophila Kinesin Light Chain PRIMARY STRUCTURE AND INTERACTION WITH KINESIN HEAVY CHAIN. Vol. 268. 1993.

37. Cross JA, Chegkazi MS, Steiner RA, Woolfson DN, Dodding MP. Fragment-linking peptide design yields a high-affinity ligand for microtubule-based transport. Cell Chem Biol. 2021 Sep 16;28(9):1347–1355.e5.

38. Antón Z, Weijman JF, Williams C, Moody ERR, Mantell J, Yip YY, et al. Molecular mechanism for kinesin-1 direct membrane recognition. Sci Adv. 2021;7(31).

39. Shubeita GT, Tran SL, Xu J, Vershinin M, Cermelli S, Cotton SL, et al. Consequences of Motor Copy Number on the Intracellular Transport of Kinesin-1-Driven Lipid Droplets. Cell. 2008;135(6).

40. Dodding MP, Mitter R, Humphries AC, Way M. A kinesin-1 binding motif in vaccinia virus that is widespread throughout the human genome. EMBO Journal. 2011;30(22).

41. Río-Bergé C, Cong Y, Reggiori F. Getting on the right track: Interactions between viruses and the cytoskeletal motor proteins. Vol. 24, Traffic. John Wiley and Sons Inc; 2023. p. 114–30.

42. DuRaine G, Wisner TW, Howard P, Johnson DC. Kinesin-1 Proteins KIF5A, -5B, and -5C Promote Anterograde Transport of Herpes Simplex Virus Enveloped Virions in Axons. J Virol. 2018;92(20).

43. Caraballo GI, Rosales R, Viettri M, Castillo JM, Cruz R, Ding S, et al. The Dengue Virus Nonstructural Protein 1 (NS1) Interacts with the Putative Epigenetic Regulator DIDO1 to Promote Flavivirus Replication in Mosquito Cells. J Virol. 2022 Jun 22;96(12).

44. Abramson J, Adler J, Dunger J, Evans R, Green T, Pritzel A, et al. Accurate structure prediction of biomolecular interactions with AlphaFold 3. Nature. 2024 Jun 13;630(8016):493–500.

45. Meng EC, Goddard TD, Pettersen EF, Couch GS, Pearson ZJ, Morris JH, et al. UCSF ChimeraX: Tools for structure building and analysis. Protein Science. 2023 Nov 1;32(11).

46. Schneider CA, Rasband WS, Eliceiri KW. NIH Image to ImageJ: 25 years of image analysis. Vol. 9, Nature Methods. 2012. p. 671–5.

47. Akey DL, Brown WC, Jose J, Kuhn RJ, Smith JL. Structure-guided insights on the role of NS1 in flavivirus infection. BioEssays. 2015 May 1;37(5):489–94.

48. Scaturro P, Cortese M, Chatel-Chaix L, Fischl W, Bartenschlager R. Dengue Virus Non-structural Protein 1 Modulates Infectious Particle Production via Interaction with the Structural Proteins. PLoS Pathog. 2015;11(11).

49. Yuan H, Luo Y, Zou J, Zhang J, Zhang J, Cao G, et al. Cellular NONO protein binds to the flavivirus replication complex and promotes positive-strand RNA synthesis. Heise MT, editor. J Virol [Internet]. 2024 Dec 17;98(12). Available from: https://journals.asm.org/doi/10.1128/jvi.00297-24

50. Laurent-Rolle M, Morrison J, Rajsbaum R, Macleod JML, Pisanelli G, Pham A, et al. The interferon signaling antagonist function of yellow fever virus NS5 protein is activated by type i interferon. Cell Host Microbe. 2014 Sep 10;16(3):314–27.

51. Mukherjee S, Singh N, Sengupta N, Fatima M, Seth P, Mahadevan A, et al. Japanese encephalitis virus induces human neural stem/progenitor cell death by elevating GRP78, PHB and hnRNPC through ER stress. Cell Death Dis. 2017;8(1).

52. Zhang M, Zheng X, Wu Y, Gan M, He A, Li Z, et al. Differential proteomics of Aedes albopictus salivary gland, midgut and C6/36 cell induced by dengue virus infection. Virology. 2013 Sep;444(1–2):109–18.

53. Samsa MM, Mondotte JA, Iglesias NG, Assunção-Miranda I, Barbosa-Lima G, Da Poian AT, et al. Dengue virus capsid protein usurps lipid droplets for viral particle formation. PLoS Pathog. 2009 Oct;5(10).

54. Heaton NS, Randall G. Dengue virus-induced autophagy regulates lipid metabolism. Cell Host Microbe. 2010 Nov 18;8(5):422–32.

55. Hafirassou ML, Meertens L, Umaña-Diaz C, Labeau A, Dejarnac O, Bonnet-Madin L, et al. A Global Interactome Map of the Dengue Virus NS1 Identifies Virus Restriction and Dependency Host Factors. Cell Rep. 2017;21(13).

56. Gao WND, Carpentier DCJ, Ewles HA, Lee SA, Smith GL. Vaccinia virus proteins A36 and F12/E2 show strong preferences for different kinesin light chain isoforms. Traffic. 2017 Aug 1;18(8):505–18.

57. Rosales Ramirez R, Ludert JE. The Dengue Virus Nonstructural Protein 1 (NS1) Is Secreted from Mosquito Cells in Association with the Intracellular Cholesterol Transporter Chaperone Caveolin Complex. 2019; Available from: 10.1128/JVI

58. Li M, Xing D, Su D, Wang D, Gao H, Lan C, et al. Transcriptome analysis of responses to dengue virus 2 infection in aedes albopictus (Skuse) c6/36 cells. Viruses. 2021 Feb 1;13(2).

59. Zhang J, Wu N, Gao N, Yan W, Sheng Z, Fan D, et al. Small G Rac1 is involved in replication cycle of dengue serotype 2 virus in EAhy926 cells via the regulation of actin cytoskeleton. Sci China Life Sci. 2016 May 1;59(5):487–94.

60. Shrivastava N, Sripada S, Kaur J, Shah PS, Cecilia D. Insights into the internalization and retrograde trafficking of dengue 2 virus in BHK-21 cells. PLoS One. 2011 Oct 4;6(10).

61. Teo CSH, Chu JJH. Cellular Vimentin Regulates Construction of Dengue Virus Replication Complexes through Interaction with NS4A Protein. J Virol. 2014 Feb 15;88(4):1897–913.

62. Bílý T, Palus M, Eyer L, Elsterová J, Vancová M, Růžek D. Electron Tomography Analysis of Tick-Borne Encephalitis Virus Infection in Human Neurons. Sci Rep. 2015 Jun 15;5.

63. Junjhon J, Pennington JG, Edwards TJ, Perera R, Lanman J, Kuhn RJ. Ultrastructural Characterization and Three-Dimensional Architecture of Replication Sites in Dengue Virus-Infected Mosquito Cells. J Virol. 2014 May;88(9):4687–97.

64. Fujimoto T, Ohsaki Y, Suzuki M, Cheng J. Imaging Lipid Droplets by Electron Microscopy. In: Methods in Cell Biology. Academic Press Inc.; 2013. p. 227–51.

65. Fruttero LL, Leyria J, Moyetta NR, Ramos FO, Settembrini BP, Canavoso LE. The fat body of the hematophagous insect, panstrongylus megistus (Hemiptera: Reduviidae): Histological features and participation of the β-chain of ATP synthase in the lipophorin-mediated lipid transfer. Journal of Insect Science. 2019 Jul 1;19(4).

66. Cortese M, Goellner S, Acosta EG, Neufeldt CJ, Oleksiuk O, Lampe M, et al. Ultrastructural Characterization of Zika Virus Replication Factories. Cell Rep. 2017 Feb 28;18(9):2113–23.

67. Joan Blanchette-Mackie E, Dwyer NK, Therese Barber L, Coxey RA, Takeda T, Rondinone CM, et al. Perilipin is located on the surface layer of intracellular lipid droplets in adipocytes. 1995.

68. Tauchi-Sato K, Ozeki S, Houjou T, Taguchi R, Fujimoto T. The surface of lipid droplets is a phospholipid monolayer with a unique fatty acid composition. Journal of Biological Chemistry. 2002 Nov 15;277(46):44507–12.

69. Shavinskaya A, Boulant S, Penin F, McLauchlan J, Bartenschlager R. The lipid droplet binding domain of hepatitis C virus core protein is a major determinant for efficient virus assembly. Journal of Biological Chemistry. 2007 Dec 21;282(51):37158–69.

70. Miyanari Y, Atsuzawa K, Usuda N, Watashi K, Hishiki T, Zayas M, et al. The lipid droplet is an important organelle for hepatitis C virus production. Nat Cell Biol. 2007 Sep;9(9):1089–97.

71. Dupont N, Chauhan S, Arko-Mensah J, Castillo EF, Masedunskas A, Weigert R, et al. Neutral lipid stores and lipase PNPLA5 contribute to autophagosome biogenesis. Current Biology. 2014 Mar 17;24(6):609–20.

72. Cui L, Mirza AH, Zhang S, Liang B, Liu P. Lipid droplets and mitochondria are anchored during brown adipocyte differentiation. Vol. 10, Protein and Cell. Higher Education Press Limited Company; 2019. p. 921–6.

73. Fontaine KA, Sanchez EL, Camarda R, Lagunoff M. Dengue Virus Induces and Requires Glycolysis for Optimal Replication. J Virol. 2015 Feb 15;89(4):2358–66.

