## Supplementary figures and images for "Kinesin light chain 1 (KLC1) interacts with NS1 and is a susceptibility factor for dengue virus infection in mosquito cells"

### Supplemental Figure 1

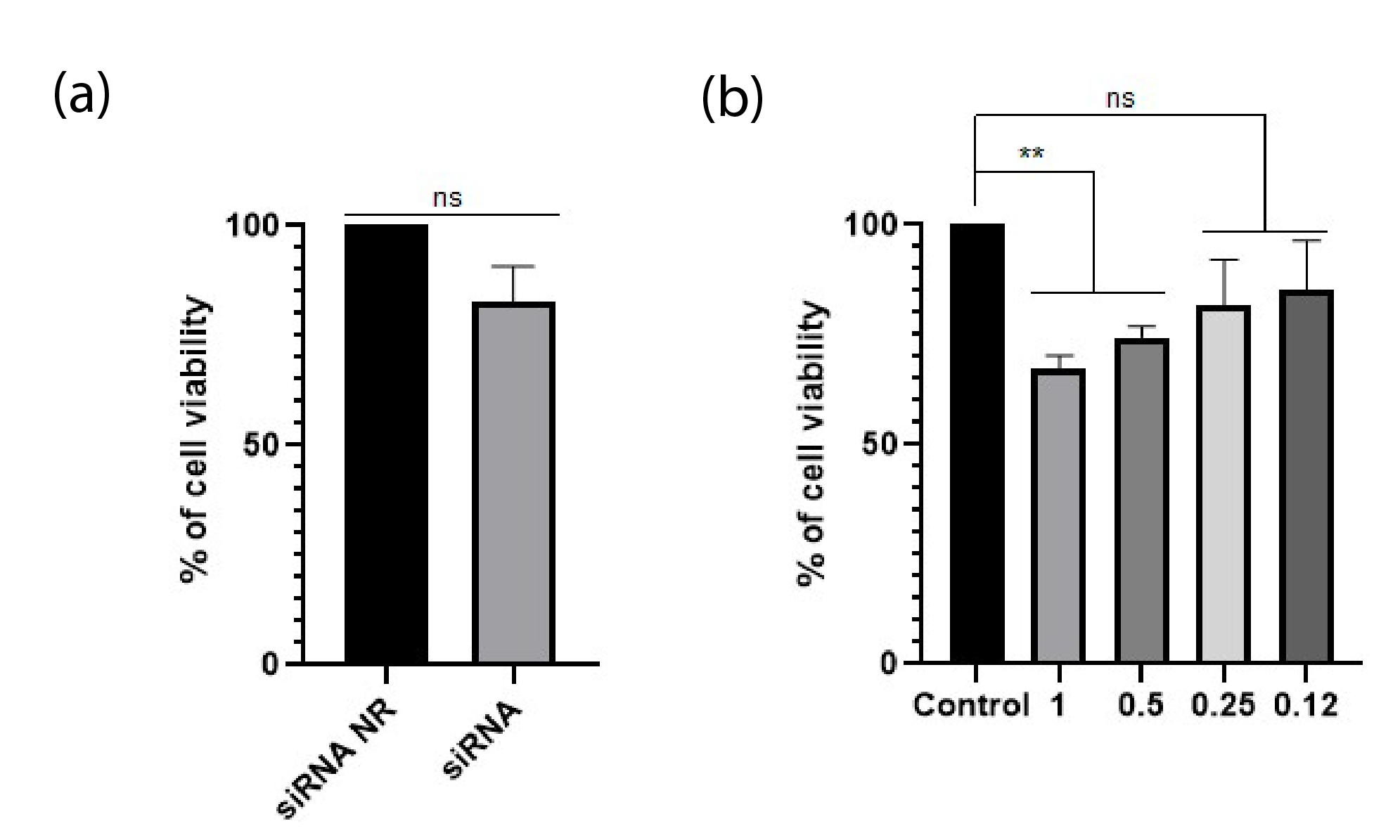

### Supplemental Figure 2

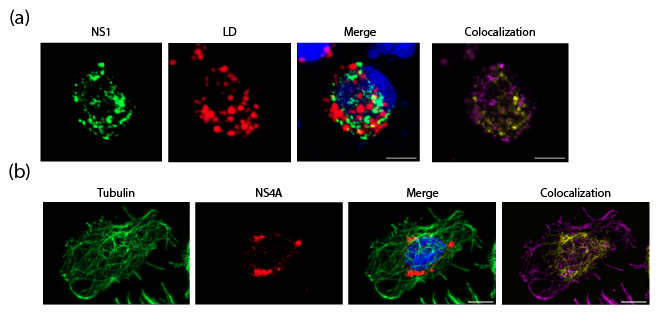
